# Three Dimensional Root CT Segmentation using Multi-Resolution Encoder-Decoder Networks

**DOI:** 10.1101/713859

**Authors:** Mohammadreza Soltaninejad, Craig J. Sturrock, Marcus Griffiths, Tony P. Pridmore, Michael P. Pound

## Abstract

We address the complex problem of reliably segmenting root structure from soil in X-ray Computed Tomography (CT) images. We utilise a deep learning approach, and propose a state-of-the-art multi-resolution architecture based on encoder-decoders. While previous work in encoder-decoders implies the use of multiple resolutions simply by downsampling and upsampling images, we make this process explicit, with branches of the network tasked separately with obtaining local high-resolution segmentation, and wider low-resolution contextual information. The complete network is a memory efficient implementation that is still able to resolve small root detail in large volumetric images. We evaluate our approach by comparing against a number of different encoder-decoder based architectures from the literature, as well as a popular existing image analysis tool designed for root CT segmentation. We show qualitatively and quantitatively that a multi-resolution approach offers substantial accuracy improvements over a both a small receptive field size in a deep network, or a larger receptive field in a shallower network. We obtain a Dice score of 0.59 compared with 0.41 for the closest competing method. We then further improve performance using an incremental learning approach, in which failures in the original network are used to generate harder negative training examples. Results of this process raise the precision of the network, and improve the Dice score to 0.66. Our proposed method requires no user interaction, is fully automatic, and identifies large and fine root material throughout the whole volume. The 3D segmented output of our method is well-connected, allowing the recovery of structured representations of root system architecture, and so may be successfully utilised in root phenotyping.

## 1 INTRODUCTION

ROOT phenotyping is the process of characterising, objectively and quantitatively, the root systems of plants [1]. It offers valuable insight into the way root systems develop, react to environmental changes and other external stimuli, and interact with their natural soil environment. Traditional approaches have involved separating the soil from the root by washing, then acquiring and analysing visible-light images [2]. This approach can offer a fairly high-throughput solution, but during the process the root structure will likely be altered, and some finer roots will be lost in the washing process: it is common for naturally 3D root structures to be flattened and imaged using flatbed scanners. Unavoidably, these destructive approaches also prevent analysis of root growth over time. Other approaches have involved growing in translucent gel, or other artificial media, preserving root structure and allowing images to be captured at multiple time points [3]. Multi-view imaging allows the 3D structure of these root systems to be recovered, but the growth and development will differ to those grown in soil and the interaction between root and soil cannot be studied where no soil is present.

Recent developments in root-soil imaging has enabled the direct examination of dynamic plant-soil interactions [4], but the data generated poses significant technological challenges. X-ray Computed Tomography (CT) is one leading technology for obtaining non-destructive root images without disturbing the root or soil structure [5]. However, a major bottleneck in the study of root systems using CT is the computational analysis of the large volumetric images produced; analysing CT data is a very time-consuming task. Automated systems have struggled to traverse complex root structure in spatially heterogeneous soil, so much of the analysis is still performed with manual or semi-manual approaches. This requires a large time investment by the user, and only becomes more difficult as the scale and throughput of modern scanners increases.

In order to accurately measure the root system, root material must be reliably segmented from soil and other objects. The complex nature of root-soil CT images makes this a particularly hard segmentation problem. CT scanners measure density, and any objects with similar density to root material, for example water, risk being misclassified. Organic material in the soil may also appear similar to roots, distracting both human and computational approaches [6]. Somewhat surprisingly, the appearance of a given root branch may vary over its length, reflecting the age of the root material, the scanning hardware and/or the distribution of material within the sample: some degree of local contextual information is required for successful segmentation. Noise distributions may also vary across a given image volume. Depending on scan resolution, roots may appear as contiguous structures, or may appear quite sparse, with only a few pixels of or no overlap between the root material regions in adjacent slices through a volume. Perhaps most challenging of all, CT volumes are typically high resolution, with root material taking up only a small proportion of the overall image. Such a difference in scale makes computationally efficient image processing at sufficient resolution difficult.

### 1.1 Root System Analysis

Many approaches to the root-soil segmentation problem of CT images have been proposed, which may be broadly classified into those that rely upon low-level image-based approaches, and those that take a more model-oriented machine learning approach. Image analysis-based methods typically use mathematical or morphological operations on the input signal, which may be 2D slices, or a 3D volume. Simple thresholding techniques have proved popular in 2D root image analysis [7], [8], but the intensity variations in volumes preclude this approach alone. In [9], the authors segment root material by identifying connected groups of pixels (connected components) via region growing. Recently, a bottom up approach to volumetric segmentation achieved good results on root volumes. Root1 [10], implemented as a plugin for the popular ImageJ tool [11], performs histogram alignment and volume stitching to create high resolution volumes from multiple scans. The volume is then enlarged to perform root segmentation at a higher-resolution than the original source. Segmentation is achieved using a multi-stage process. Pore space is removed by detecting air/soil boundaries using an edge detection algorithm, a step that makes thresholding the image more straightforward. A bilevel user-defined threshold is used to separate out root material from the remaining background. Morphological operations and other image filters are applied to remove as much noise as possible, before the root system is extracted as a user-selected connected component in the separate Volume Graphics (Volume Graphics Gmbh, Germany) software package. Root1 can offer competitive results on some volumes, and is resilient to changes in species and soil. However, it is susceptible to noise, and produces many false positives that must be removed by a user via a separate application. The threshold is also user defined, meaning that the pipeline of loading, analysing and moving volumes between packages is time consuming.

A tracking based segmentation method was proposed in [6], called RooTrak, which uses level-sets [12] to track the root as it progresses downward through the stack, providing some contextual information. The level-set method evolves a front that represents the boundary between root and soil. The function evolves outward and inward, fitting to the image data, while preserving constraints on shape and curvature. RooTrak employs additional techniques, such as back tracking upward through the stack, to traverse the maximum possible extent of the root system. RooTrak requires the selection of a single seed point by the user, and a few parameters that define the expansion of the front, but this is comparatively low interaction compared with the bottom-up approaches discussed above. However, image noise, finer root detail and the edges of the container will often cause tracking failure, which require manual reinitialisation by the user. In practice, the use of RooTrak may involve substantial user intervention on challenging data.

### 1.2 Image Segmentation with Deep Learning

In recent years, Deep Convolutional Neural Networks (CNNs) have established themselves as a dominant technique in Computer Vision. CNNs have also shown stateof-the art results in semantic segmentation (pixel-level prediction) [13]. The fully convolutional network (FCN) [14] was one of the first networks introduced for pixel-wise semantic segmentation which could be trained end-to-end. The typical fully-connected layers that perform classification were replaced by 1×1 window filters, allowing the network to produce 2D output. However, at this early stage segmentation accuracy was coarse, with ill-defined boundaries and poor performance on small objects. More recent methods have tackled this problem by attempting to preserve more information while an image passes through the network. Since the loss of information happens during image down-sampling, a common component of deep networks, work has proposed replacing these with dilated convolutions [15], [16], [17]. Dilated convolutions help enlarge the receptive field of the filters in a way that larger context may be considered, without increasing network complexity. In [18], deformable convolutions were used, in which the filters field of view is determined by using the input features. SegNet [19] proposed using unpooling layers followed by convolution with learnable filters to produce denser feature maps, and a higher resolution output. DeepLab [20] proposed using atrous convolution for dense prediction along with increasing the input receptive field and fully connected conditional random fields (CRF). Other works have proposed using deconvolution modules during the upsampling pipeline [21], [22].

Regardless of the individual layers used, in the majority of cases it has become commonplace to separate the convolution and downsampling layers of a network (the encoder) with the upsampling or deconvolutional layers (the decoder). Encoder-decoder networks have a symmetric structure, in which deconvolutional or unpooling layers have identical resolution to the corresponding convolutional and pooling layers [23], [19], [24]. Features are passed through the network as normal, but also skip from the encoder to the decoder, providing low level image information at the stage in the network where the output is being produced. Skip layers copy the features from the down sampling path to the corresponding resolution layer in the upsampling path, combining them with either a sum or concatenation operation. Some methods proposed stacks of encoder-decoders for end-to-end segmentation [25], [13], [26]. The stacked hourglass network [25] consists of series of encoder-decoders with residual blocks that incorporate residual features alongside with the spatial information for learning [26], [27]. Residual blocks have been widely adopted in modern CNN design, allowing deeper networks to be trained more quickly. They have underpinned state-of-the-art performance in large scale image recognition [28], and semantic segmentation [29]. The stacked hourglass network further improved on encoder-decoder architectures by performing additional learning within the skip connections, controlling what information flows from shallow layers to deeper layers of the network. Fu et al. [13] proposed a stacked deconvolutional network in which intra and inter connection skip layers were used to improve the flow of information and backpropagation with hierarchical supervision. Their method achieved high performance results with a lower network complexity and size.

### 1.3 Volumetric Segmentation

U-Net is an encoder-decoder structure that was originally proposed for medical image segmentation [30]. Within the field of root image analysis, Smith et al. [31] proposed using U-Net for segmentation of roots from soil in 2D rhizotron images, comparing their method to the Frangi vesselness [32] filter, an image-based approach originally designed for segmentation of (e.g. blood) vessels that have similar structure to roots. U-Net was extended in [33] into three dimensional convolutional and pooling layers for voxel-wise volumetric segmentation. V-Net [34], represents similar work using a 3D fully convolutional encoder-decoder designed for volumetric semantic segmentation. In the medical domain, volumetric networks have become commonplace for the segmentation of MRI images [35]. These tasks are made somewhat easier by the limited resolution typically used in medical MRI (and CT) imaging. Root CT imaging produces volumes sometimes orders of magnitude larger. This is made more challenging by the fact that the root system may span a wide area, but with individual fine roots only spanning a few voxels. This contrast means that common techniques to tile or downsample volumes for efficiency are, as we shall show, ineffective in this domain.

### 1.4 Contribution

In this paper we propose a segmentation method for volumetric segmentation of root systems from soil in CT. We use a volumetric deep network with an encoder-decoder architecture, but with significant architectural changes to directly address the shortcomings of deep learning on large volumes. Rather than a compromise between a sufficiently large receptive field and a high-resolution input, we propose a two branch network, one that examines a high resolution volume with a small receptive field, another that examines a lower resolution volume but with increased input size. The branches are then combined into an output that draws from the higher resolution and wider context these branches provide. We train the network with multiple auxiliary loss functions to incorporate both semantic and local features in each branch. The network is trained using a dataset of very large CT images captured with a Phoenix v|tome|x m micro CT scanner. The images were manually segmented by an expert and used as ground truth for training and evaluation of the network. We perform extensive quantitative analysis of segmentation accuracy, showing that our proposed approach offers higher accuracy over both existing root segmentation tools, and other network architectures commonly used for semantic segmentation. We also demonstrate that incremental learning using hard negative examples is able to further increase performance. Our approach requires no user interaction or parameter selection in order to segment unseen volumes, offering a crucial step towards full automation of root CT. The major contributions of this paper are:

- We develop and train a new network architecture comprising a multi-resolution parallel pipeline of encoder-decoders for the volumetric segmentation of root systems. The network is trained end-to-end, and incorporates a high-resolution small receptive field, with a lower-resolution large receptive field. This compromise improves the resolution of the network on fine root detail in very large image volumes. To our knowledge this is the first application of volumetric CNNs to the problem of plant root segmentation in CT images.
- We utilise a multi-loss training approach that distributes the learning process to each pipeline individually, forcing each parallel branch to learn different tasks to contribute to the overall segmentation accuracy. We evaluate this multi-loss against the same network with a single loss.
- We evaluate our proposed network architecture against state-of-the-art networks in semantic segmentation, all adapted for volumetric segmentation. We show improved performance on all metrics.

The rest of this paper may be summarised as follows: Section 2 outlines in more detail the core concepts of encoder-decoder networks, and our parallel network design for volumetric segmentation. We then describe the dataset used for training and testing, and the training procedure. Section 3 describes the metrics we use for quantitative comparison, and provides experimental results against existing root segmentation techniques and state-of-the-art network architectures in segmentation. Section 4 discusses the advantages and limitations of the proposed method and its application to root system analysis. Finally, section 5 presents a conclusion and possible directions for future work.

## 2 METHODS

This section describes our proposed approach to root segmentation in volumetric images. We first outline two approaches to tackling the problem of high-resolution CT segmentation by either downsampling the volume for efficiency, or considering only very small tiles of the original data. We then outline the multi-resolution network and loss functions for joint prediction.

### 2.1 Native Resolution Encoder-decoders

Encoder-decoder networks have become a popular approach to semantic segmentation. Popular examples include FCN [14], Segnet [19], U-Net [30], and Stacked hourglass [25]. The base of our network uses an architecture similar to a stacked hourglass, with modifications to handle volumetric data. The architecture of this network is presented in Figure 1.

**Figure 1:**
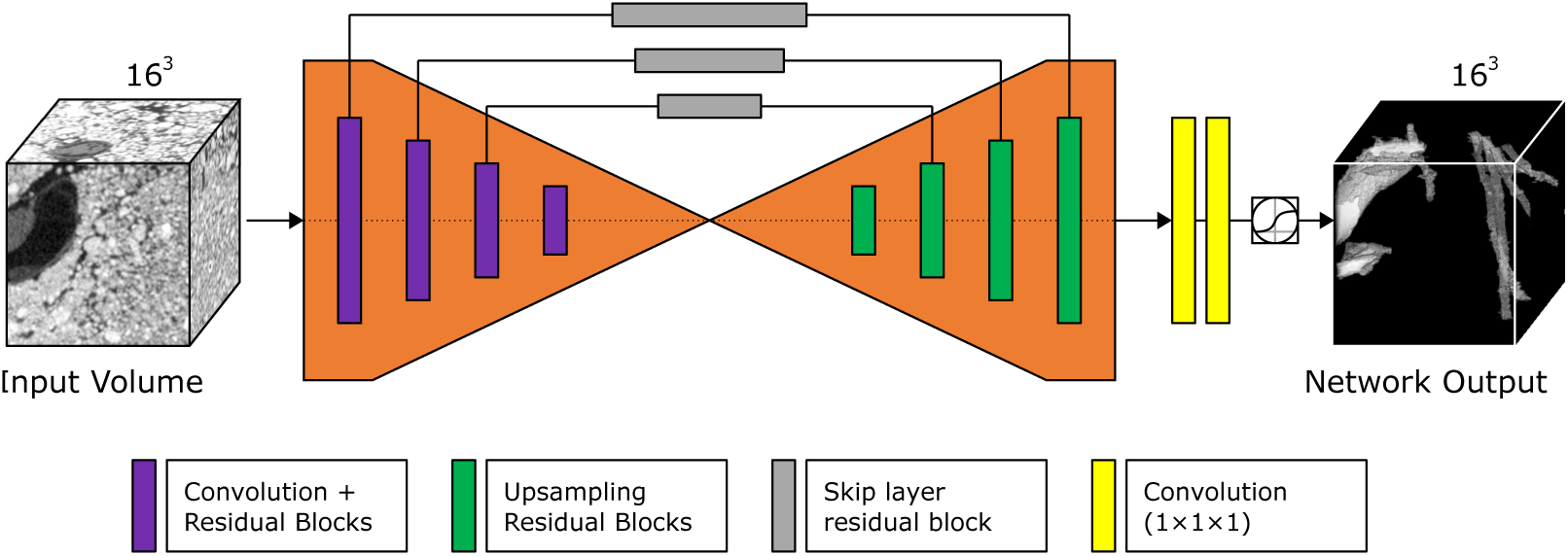
The architecture of a native resolution encoder-decoder. The network performs segmentation by using filtering and downsampling operations to encode an input into a feature space, before upsampling this back to the original resolution.

Unlike a stacked hourglass network, we modify the architecture from a 2D spatial network into a 3D volumetric network through the replacement of layers as appropriate. We limit the input size to 163 pixels, and use a feature size of 128 throughout the network in order to improve efficiency when processing volumes. The output of the network is generated by two 1×1×1 convolutional layers, which draw on features generated within the network to produce a volumetric segmentation output. During evaluation of this network in isolation we use a stack of two encoder-decoders end-to-end. When used as a component in a larger network, we use only a single encoder-decoder.

### 2.2 Larger Fields of View

Volumetric networks such as that in section 2.1 are memory intensive, having essentially an extra dimension above traditional CNNs. The network above will only accept small volume inputs of approximately 163 before memory consumption becomes prohibitive, and further reductions in depth or feature size would be necessary. While individual roots may be smaller than this input size, the limited field of view makes it challenging to discern what is root material, and what is other soil matter. We develop a second network that incorporates additional downsampling prior to the encoder-decoder, shown in Figure 2. This architecture severely constrains the resolution for the majority of the network, allowing a larger input FOV of 128^3^ pixels.

**Figure 2:**
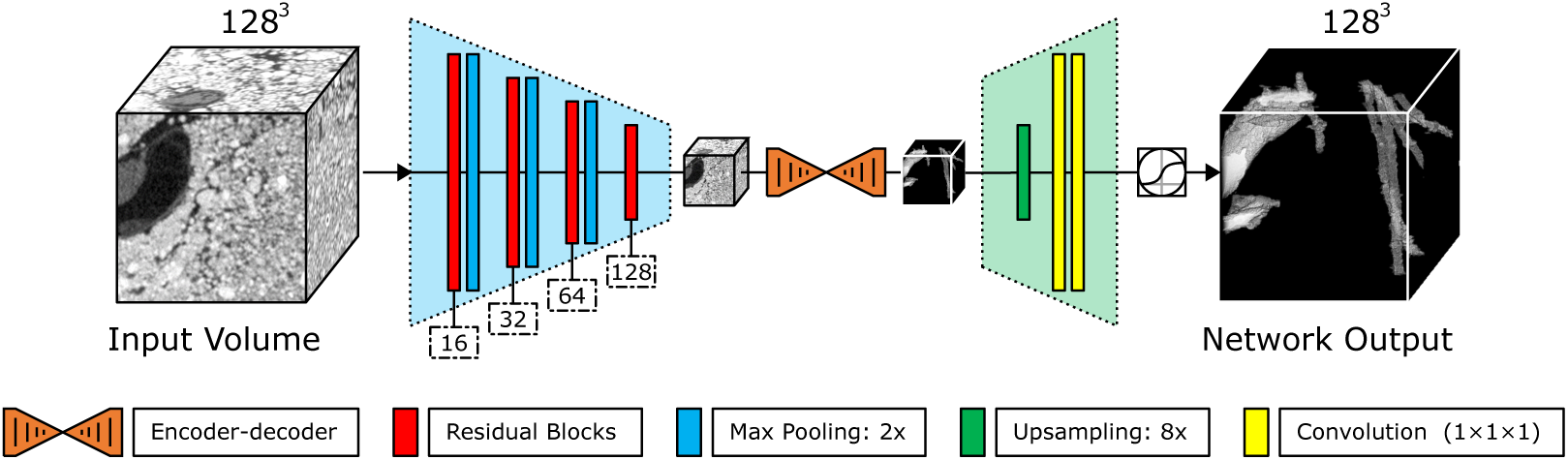
The architecture of an encoder-decoder that uses additional downsampling to operate at lower resolution. This allows it to accept larger input volumes, and so wider fields of view.

**Figure 3:**
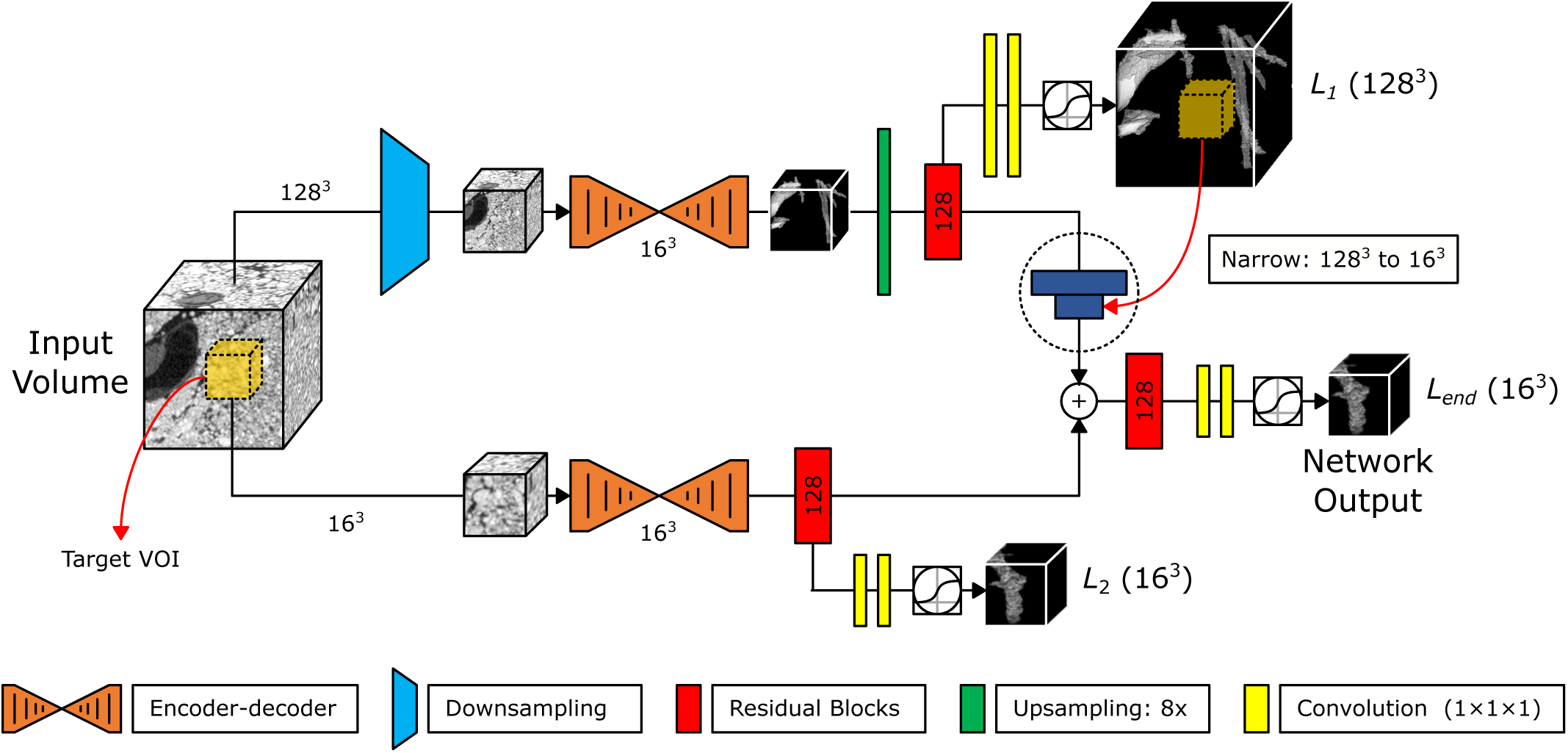
The architecture of our proposed multi-resolution approach. The network uses two paths, a native resolution path accepting smaller volumetric input, and a downsampled path accepting a wider field of view. The paths are spatially aligned and concatenated prior to segmentation. The network is trained end-to-end, and utilises auxilliary loss funtions on each path.

We incorporate residual blocks during the initial downsampling process, increasing the feature size progressively up to 128 features before these are passed into the encoder-decoder. Following the encoder-decoder, we upsample the features back into native resolution, before 1 × 1 × 1 convolutional layers are used to provide segmentation output. This architecture provides a much larger FOV than the smaller network, at the cost that the additional downsampling provides a lower working resolution for segmentation. For either of these networks, successful segmentation will be a compromise between an acceptable resolution, and an adequate field of view.

### 2.3 Multi-resolution Encoder-decoder Architecture

In order to utilise both local pixel information and the wider contextual information from the surrounding FOV, we propose a parallel architecture that operates at multiple resolutions. This network utilises two parallel pipelines, each considering a different input volume size. The outputs of these paths are combined, with final segmentation utilising features from both. The structure of the proposed network is presented in Figure 2.

The upper path consists of a downsampled encoder-decoder network, which takes a larger 1283 volume as input and extracts features of the root system at a coarse resolution. The lower path is a conventional encoder-decoder, taking the centre of the volume at native resolution and producing finer segmentation detail for that patch. We refer to these paths as the downsampled and native paths respectively. Since both paths represent different fields of view, the features of the downsampled path are cropped after upsampling to align properly with the native path, before a final prediction is made using both sets of concatenated features. Two 1 × 1 × 1 convolutional layers provide the final prediction from this feature space.

The network is trained using stochastic gradient descent with a Binary Cross Entropy loss function. We first train the network end-to-end using a single loss function *L*_*end*_. We refer to this network as the single loss multi-resolution (SLMR) network. While using the final heatmap alone as a mask from which to calculate the loss is convenient, it may produce a bias towards the weights learned in the native path of the network. Since the outer FOV is cropped prior to applying the loss, the outermost feature activations are not used during back propagation. While they may have some effect within the convolutional layers, this is not made explicit. We apply two additional losses, *L*_1_ and *L*_2_ to each path in order to learn optimal segmentation within both branches. The total loss function criterion is then calculated using joint prediction of all three losses, in what we call the multi-loss multi-resolution (MLMR) network. It should be noted that the same smaller ground truth is applied at both *L*_1_ and *L*_*end*_, whilst the larger FOV ground truth is applied at *L*_2_. All three losses ensure that the network utilises both local pixel information and wider FOV in performing segmentation, a property we have found to improve segmentation accuracy.

## 3 RESULTS

We evaluate our method on volumetric CT images of intact wheat roots grown in soil captured at the University of Nottingham’s Hounsfield Facility. Each volume was annotated by an expert using a manual, region growing segmentation approach within the Volume Graphics software package. This provides ground truth. We provide a comparison against an existing bottom-up tool in this domain (Root1) along with a number of state-of-the-art network architectures in order to investigate the performance of our multiresolution network. This section first describes the data and ground truth production, the network setup and training configuration, and comparative experimental results.

### 3.1 Data

We use a dataset of 47 volumetric images of intact wheat roots in soil captured using a Phoenix v|tome|x m micro CT scanner. 10-day-old wheat plants were grown in a sandy loam sand mixture and CT scanned at a resolution 54*µm* per voxel in each axis. The voxel resolution of the images varies from 1626 to 1720, and 1633 to 1706 pixels in the X and Y dimensions respectively, with the average of 1670 and 1667 pixels. The number of slices (Z-depth) ranges from 2187 to 2850 pixels with the average of 2541 pixels and slice thickness of 100 nm. This dataset includes roots of six different lines of wheat. The images were saved as stack using 8 bits per pixel precision, and randomly split into training, validation and testing sets of 32, 8 and 7 volumes respectively.

### 3.2 Training Process

The network is implemented using Torch7 [36], and trained on an Nvidia Titan X with 12 GB RAM. The number of features at each layer for the internal encoder-decoders is set to 128, except where noted otherwise in Sections 2.2 and 2.3. During training, mini-batches of 5 × 128^3^ crops were obtained from volumes at random. While the volumetric images are very large, only a small proportion of voxels contain root material. This means that the majority of random volume positions containing no root material, so do not always provide helpful training data. This imbalance between the foreground and background leads to challenges when training these networks on random mini-batches, where high accuracy may be achieved simply by predicting no root material at all. Our approach is to produce crops nearby known root locations from the ground truth a proportion of the time, ensuring that a percentage of each batch contains root material. Prior to training, a random sampling of coordinates containing root material are drawn from each volume. During training, whenever a volume is used, a position nearby one of these pre-sampled points is chosen 30% of the time, with the remaining patches cropped entirely at random, ensuring only that the crop remains within the bounds of the image. We applied the same sampling regimen during validation to ensure that challenging crops were chosen.

Each network was trained using rsmprop [37] with a learning rate 2.5 × 10^*-*4^. The batch size and/or parameters of each network were adjusted as appropriate to maximise their utilisation of the available GPU memory, to ensure a fair test. Each network was trained for a maximum of 100 epochs, and the epoch with the highest validation performance was kept. The final predications are obtained by thresholding the output of the final layer, with every location segmented during testing. Figure 4 plots the training and validation performance for the different network architectures. The loss value of the training set is calculated at each epoch, with the validation loss calculated every 5 epochs.

**Figure 4:**
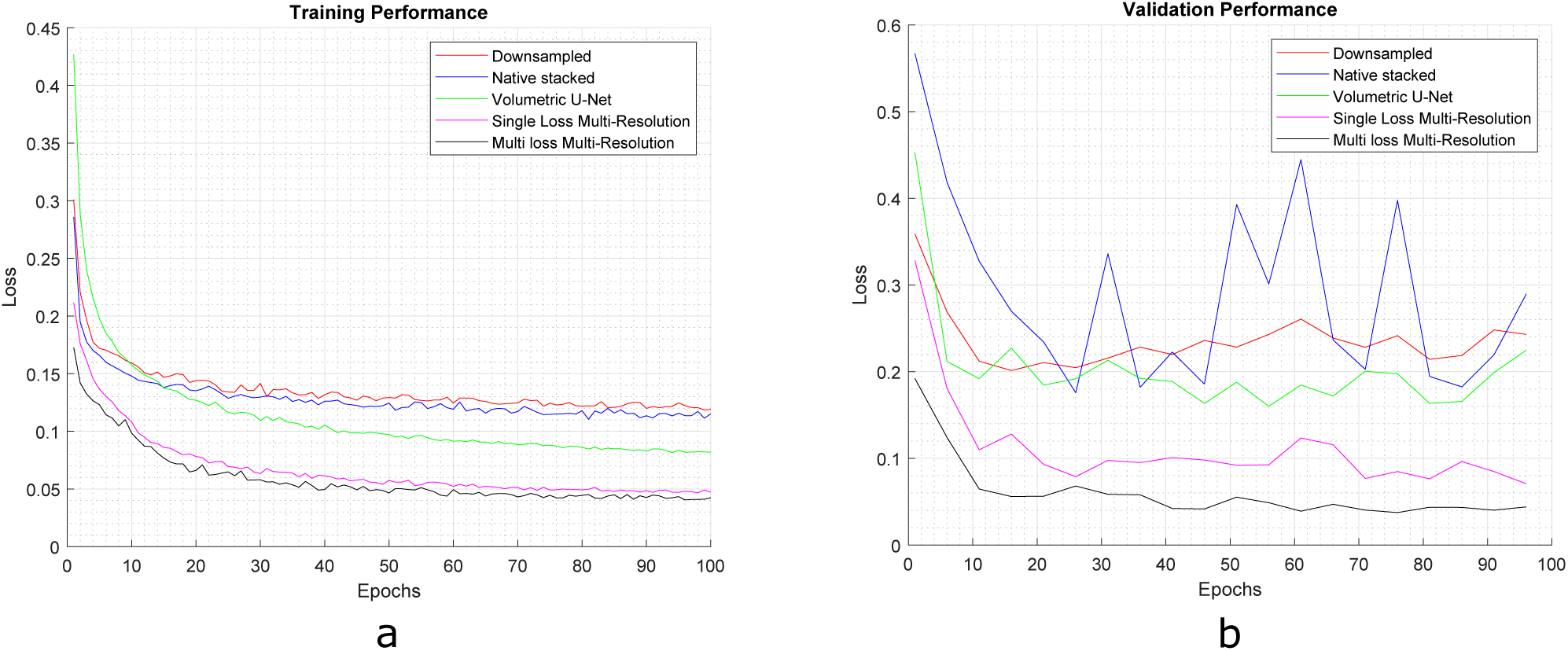
The loss calculated for training and validation per epoch for different architectures of the learning based methods.

We found that the both variants of the multi-resolution network converged faster than the other architectures, and the multi-loss multi-resolution network provided the most stable validation performance.

### 3.3 Evaluation Metrics

During training we evaluate the training performance of the segmentation by recording successes (true positives and true negatives) and errors (false positives and false negatives) on a per-voxel basis. We calculate these metrics across all minibatches for an epoch, which based on our training sampling regime includes the sampling bias towards selecting regions containing root material over those chosen at random. During validation, we use two passes and combine the results to ensure that a sufficient sampling is obtained from each volume. Once training is complete, each network is finally evaluated by processing the entire stack at every location, producing an output segmentation that may be compared to the ground truth.

We evaluate our network using a number of commonly used metrics. Since root voxels are greatly outnumbered by background, we do not consider measures dependent on true negative success (such as overall percentage accuracy), as these values are often close to 1, and less helpful. We first calculate precision and recall, useful for identifying general segmentation accuracy, but also distinguishing between under- and over-segmentation. These are calculated as:

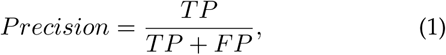

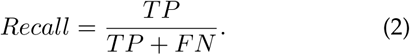

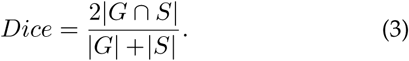

It is also common to calculate Dice from precision and recall, where it is referred to as *F*_1_ score, as follows:

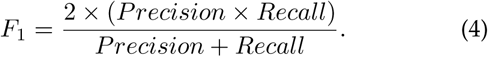

We present only Dice here, as these measures are equivalent. Dice scores range from 0 to 1, with values closer to 1 indicating better segmentation and closer adherence to the ground truth.

We also calculate Jaccard index, or Intersection over Union (IoU). This measures the overlap between the prediction and ground truth, and is calculated as:

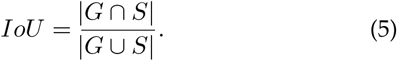

Similar to Dice, IoU scores closer to 1 indicate better segmentation performance.

### 3.4 Quantitative Evaluation

Figure 5 provides an overview of the networks discussed in section 2, including the multi-resolution network we present as the contribution of this work. We compare the accuracy of native and downsampled encoder-decoders, and two variants of our multi-resolution network, one with auxiliary loss functions applied to each branch.

**Figure 5:**
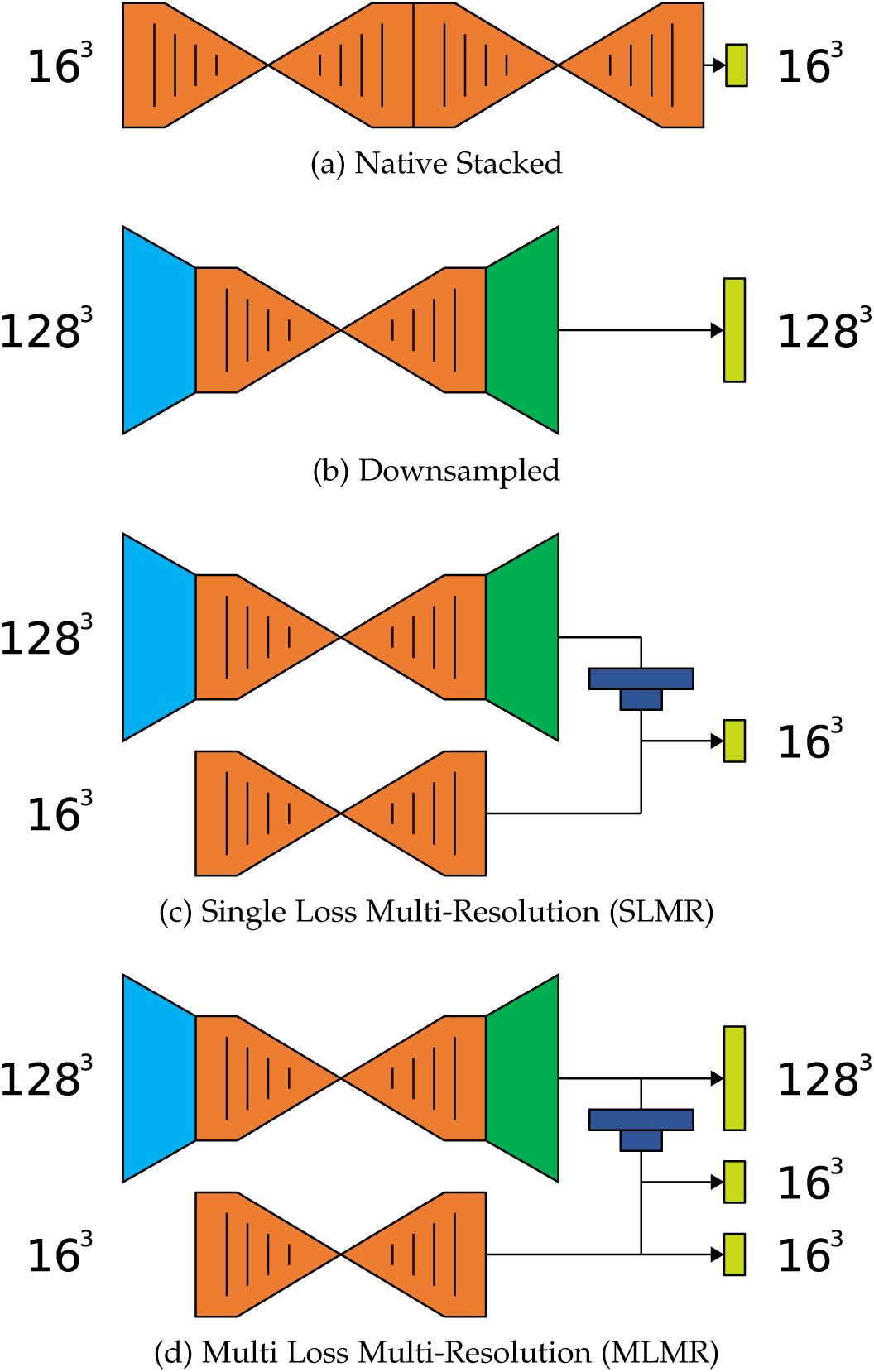
Different volumetric hourglass based network structures and the corresponding details for architectures that can be implemented on a 12GB GPU.

In order to provide a reliable comparison with existing techniques, we also evaluate a volumetric U-Net [33] as a baseline architecture, and an image processing pipeline driven by the Root1 software package. Root1 performs segmentation using edge detection, filtering and thresholding, and forms part of a pipeline incorporating dilation and connected components typically completed within the separate Volume Graphics application. Volumes were first segmented using Root1, stacks were saved and then loaded into Volume Graphics. Optional dilation of varying levels was applied in order to connect roots that may be separated by small areas of background, before user-selected seed location(s) and the 3D region growing tool were used to select the root system as a contiguous connected component. It is worth noting that this semi-automatic noise removal pipeline could also be applied to the output of any of the deep networks considered here, however we wanted focus on full automation of these machine learning techniques. Nevertheless, we utilise the entire process for Root1 and present these results in order to provide a comparison between our approach and a leading existing pipeline in full. As noted below, we also remove any segmented soil container from the U-Net and native networks specifically, however we perform no noise removal or post processing on our multi-resolution networks. Table 1 presents a comparison of results across all validated approaches.

**Table 1:** Segmentation accuracy metrics for the MLMR network when trained both using our original data augmentation approach, and following incremental learning using hard negative examples.

| Method | Precision | Recall | Dice | IoU |
| --- | --- | --- | --- | --- |
| Image based approaches |  |  |  |  |
| Root1 - no dilation + cc | **0.764** | 0.477 | 0.579 | 0.414 |
| Root1 - dilation x2 + cc | 0.319 | 0.786 | 0.445 | 0.291 |
| Root1 - dilation x3 + cc | 0.209 | 0.865 | 0.325 | 0.199 |
| Deep learning based approaches |  |  |  |  |
| Volumetric U-net | 0.453 | 0.589 | 0.508 | 0.342 |
| Native stacked | 0.330 | **0.775** | 0.461 | 0.301 |
| Downsampling | 0.646 | 0.323 | 0.428 | 0.274 |
| Single-loss multi-resolution | 0.673 | 0.767 | 0.717 | 0.559 |
| Multi-loss multi-resolution | 0.733 | 0.750 | **0.740** | **0.588** |

The results in Table 1 show that deep learning approaches often offer improved performance over traditional imaging pipelines, despite human intervention in the latter. Root1 without dilation but including manual noise removal outperformed the common deep network architectures on combined metrics. Our multi-resolution approach outperforms all other deep learning techniques by a margin of 0.246 IoU. The results also show that, depending on the approach, there is often a trade-off between high precision, and high recall.

We used three alternate pipelines for Root1 software, all derived from the outline in the original work. We segmented each root volume in the test set, and then applied varied levels of dilation to the output. Finally, a user selected one or more seed locations to extract connected components. The pipeline without dilation showed a competitive precision score. Manually selecting connected components means that much of the noise is removed, but also any roots not connected to the main root structure. This is borne out in the precision score of 0.764 and yet a low recall of 0.477, indicating half the root system is missed on average. Introducing dilation substantially increases recall, as roots that are only slightly separated may join the main root mass and be included in the final segmentation. The drawback of this approach is that the root system is now oversegmented, for example a precision of 0.209 indicating only 20% of the marked root pixels are true positives. We found it hard to find an adequate balance between precision and recall using this technique, and correction of segmentation error placed a heavy burden on the user to find root material.

The conventional stacked encoder-decoder network achieved the best recall score of 0.775, but its precision of 0.330 is lower than the average, and represents a very high rate of false positives. It obtains among the lowest segmentation overlap, i.e. Dice score of 0.301 and UoI of 0.461. In essence, the network is prone to over-segmentation; it avoids missing roots, but provides too much noise and misclassified background. The volumetric U-Net shows increased precision of 0.453, indicating fewer false positives when compared to the native network. The U-Net consumers fewer resources, meaning input volumes of 64^3^ are possible, compared with 16^3^ for the while for native network. This represents a wider field of view for the network, incorporating more root material, and increasing a networks ability to distinguish between true root material and clutter. Despite this, the recall for the network drops to 0.589, suggesting that a shallower network that does not incorporate residual blocks is less able to discern root material from background. It should also be noted that in some volumes the volumetric U-Net and native resolution networks incorrectly segment the container holding the soil as root material. For these two methods, this region was manually eliminated from the output segmentation mask to establish a fair comparison in the soil region with other methods. No post-processing was performed on the output of either multi-resolution method.

Nevertheless, the wider field of view utilised by the U-Net appears to benefit precision, and the downsampling network introduced in 2.2 was designed to increase further the receptive field size of the native network to 128^3^. Indeed, results show that the increased field of view improves the precision to 0.646, representing a lower false positive rate. However, the recall has been reduced significantly to 0.323, indicating that many areas of root material have been omitted. This is unsurprising, a lower resolution network discards much spatial information in favour of a wider field of view, lateral roots in particular are small structures, and many are under-segmented by this network. The higher precision metric does not indicate a better performance overall, with the downsampling network achieving a Dice score of 0.428, and UoI 0.274.

We hypothesised that a network incorporating both a wider field of view, and a native resolution path, would out-perform methods that selected only one of these approaches; in Section 2.3 we proposed such a multi-resolution architecture. The single loss strategy combines the two parallel networks achieved precision and recall values of 0.673 and 0.767 respectively. On combined metrics, Dice and UoI increase to 0.717 and 0.559, outperforming the previously discussed methods. The precision of this network remains similar to that of the downsampling approach, but with substantially increased recall over that network, a fact we attribute to the included native resolution path. Adding auxiliary loss functions allows us to train each path independently, enforcing the role of each path, and further increasing performance. The MLMR network shows increased precision of 0.733 and recall of 0.750. It is worth noting that while the native network and Root1 + no dilation score slightly higher in individual metrics, they achieve this with a substantial loss in recall and precision respectively. The MLRL network avoids this trade-off, achieving a Dice score of 0.740 and UoI of 0.588 across the test set.

Figures 6 and 7 show sample outputs from each of the tested approaches and provide a qualitative overview of the relative performance and modes of failure. Figure 6 presents results on sample slices of a volume, ranging from the entire soil column through to highly zoomed sections containing one root. The results confirm our quantitative analysis, showing that networks operating at native resolution offer improved recall — they find more roots — and those applying downsampling offer improved precision — they do not oversegment. However, in all but the multi-resolution networks, high performance in either of these comes at the cost of low performance in the other.

**Figure 6:**
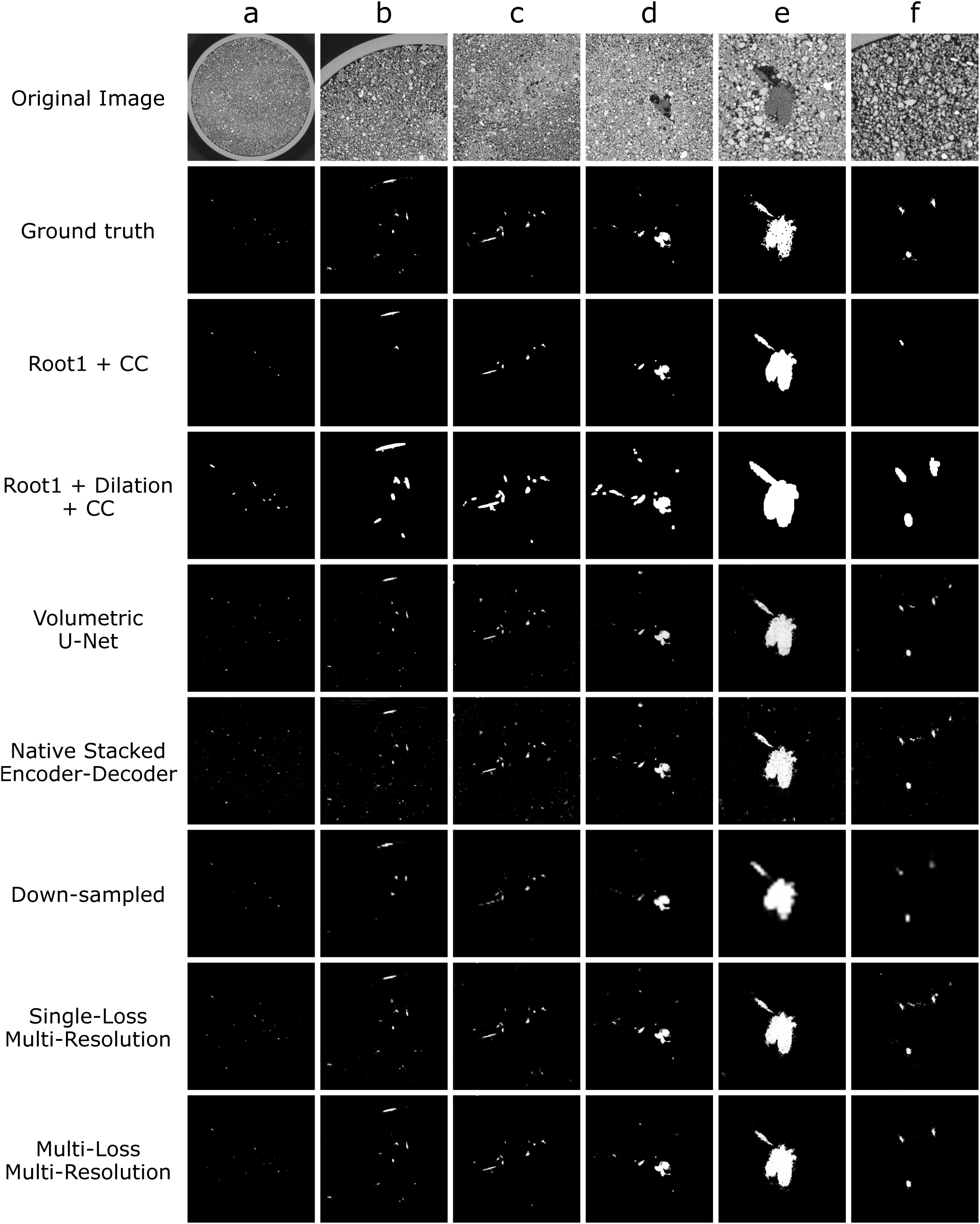
The segmentation results in a CT slice: original CT image, manual segmentation (ground truth), Root1 and connected component, Root1 and dilation and connected component, volumetric U-Net, conventional stacked hourglass, downsampling hourglass, single-loss multi-resolution hourglass, multi-loss multi-resolution hourglass. a-f) different resolution from different slices.

**Figure 7:**
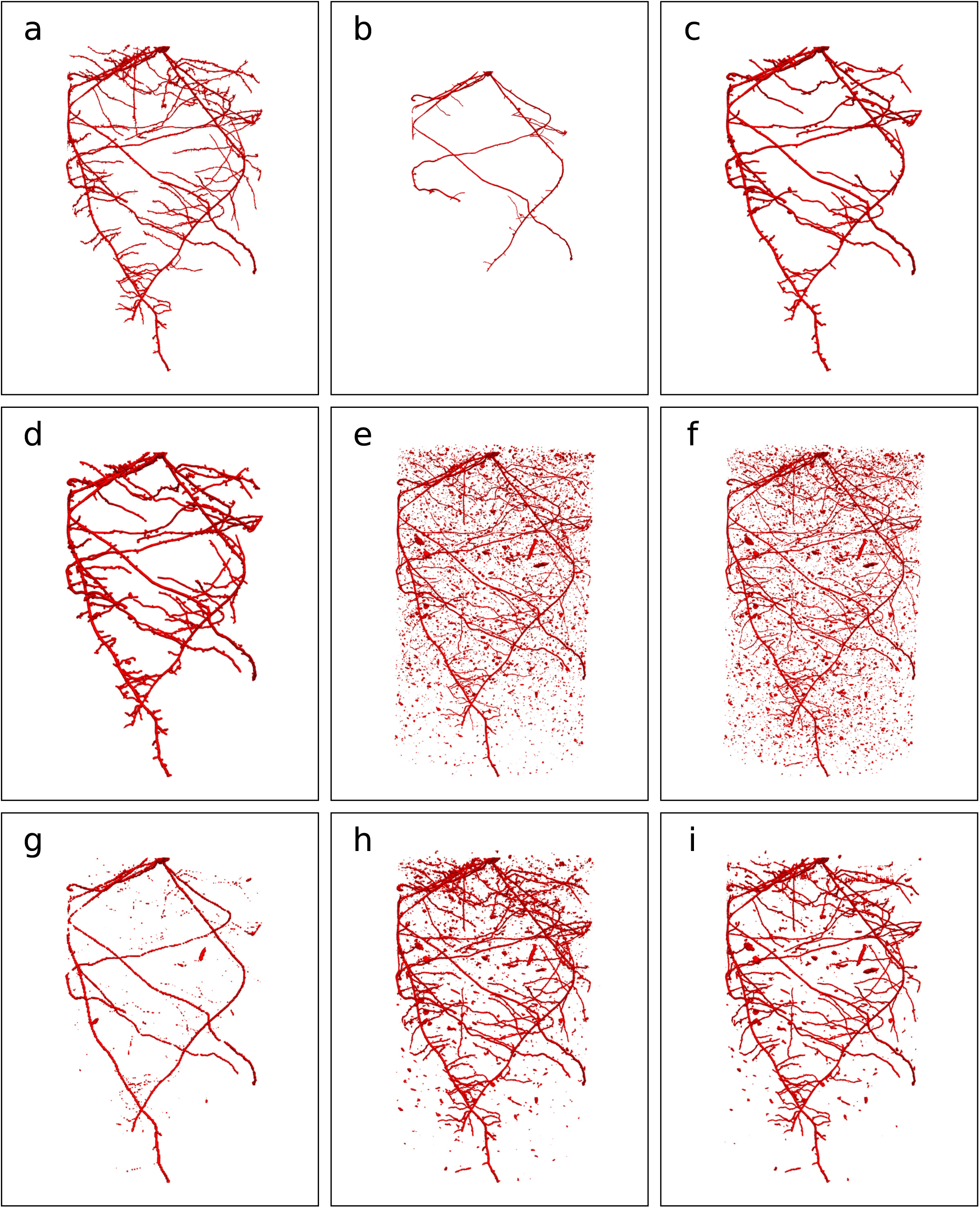
Volume rendered from the segmentation masks: (a) manual segmentation (ground truth), (b) Root1 and connected components, (c) Root1 + dilation x2 and connected components, (d) Root1 + dilation x3 and connected components, (e) volumetric U-Net, (f) native stacked encoder-decoder. (g) downsampling network, (h) single-loss multi-resolution hourglass, Ii) multi-loss multi-resolution hourglass

The volumetric U-Net was able to localise and segment much of the root material, however it also over-segments, producing a large number of false positives throughout the volume. The native stacked encoder decoder is able to discern finer detail, producing more accurate segmentation of individual roots, but as with the U-Net it produced a great many false positives. These typically occur on particles soil components that appear similar in density to root material, and where a small field of view is unable to discount them based on shape. By extending the input FOV and taking structural features of the roots into account, the downsampling network produced many fewer false positives. However, many finer roots such as laterals were not segmented, as can be seen in Figure 6 (Down-sampled row) and Figure 7f. The fidelity of the root boundaries is also reduced by the drop in resolution. Our major contribution in this paper is two approaches to a multi-resolution network, utilising different loss function strategies. Both the multiresolution single-loss SLMR and multi-loss MLMR architectures successfully produce finer segmentation detail that is quantitatively close to the ground truth. The single-loss architecture produces a higher number of false positives, more clearly shown in Figure 7h. This is due to the loss function only being applied to smaller 16^3^ patches, and not calculating a loss over the wider FOV in the downsampling branch. A multi-loss approach considers this important contextual information in the surrounding root structure, successfully segmenting fine root boundaries while also suppressing many of the false positives. Some false positives remain, we address these in 3.5 using an extended training approach.

Root1 was able to segment much of the major root system, but omitted some finer root detail. Due to the use of a connected component approach to noise removal, Root1 does not produce false positives where they are disconnected from the root system. However, this also causes roots to be omitted from the output if they have even minor disconnections, causing many roots to be lost (Figure 7,b). This is also not an automatic process, with this root system requiring multiple user-located seed positions for the connected component process. We found a single seed location in Figure 7,b did not produce satisfactory results, and this figure was generated using three seed locations. Adding dilation to the post-processing of Root1 allows the recovery of smaller roots, at the cost that the root boundary no-longer aligns with the ground truth, making measures such as root diameter and volume unreliable. Some finer root detail was still lost after dilation, such as the vertical root near the top of the volume in Figure 7.

### 3.5 Incremental Learning using Hard Negative Examples

While the MLMR network produced fewer false positives than other networks that incorporate native resolution data, some false positives remain, for example those shown in Fig. 7.f. One solution to this problem would be to eliminate small isolated groups of voxels using a connected component algorithm, similar to that used in the Root1 pipeline. Mathematical morphology is also commonly used in segmented root images to improve connectivity and reduce noise. This approach is not robust; the amount of dilation or erosion depends on the dataset and may even change depending on the individual image. Successful segmentation will often require a human-in-the-loop approach to postprocessing. Our approach remains fully automatic: we wish to remove noise at a network level.

We use the false positives identified in the training set as additional hard negative examples when incrementally training the network. A snapshot of the MLMR network that poor precision was used to completely segment the training dataset, predictions were compared to the ground truth and the false positive regions were extracted and combined with the list of available training patches used in section 3.2. In this new training regime, the 30% of minibatch samples specifically targeted at root material were still used, along with an additional 20% from the hard negative, i.e. false positive, locations. The remaining 50% of points were still chosen at random from any location in a volume. This approach ensures that there is a balance between sampling the seldom seen but important positive and negative examples, along with samples of random image regions. The network was then trained for an additional 100 epochs as per our original training process, and the best validation performance was selected. Table 2 shows a quantitative comparison of the segmentation results for the best performing MLMR network, and the refined model trained with additional hard negative examples.

**Table 2:** Segmentation accuracy metrics for the MLMR network when trained both using our original data augmentation approach, and following incremental learning using hard negative examples.

| MLMR Network | Precision | Recall | Dice | UoI |
| --- | --- | --- | --- | --- |
| Incremental learning | 0.733 | **0.750** | 0.740 | 0.588 |
| No incremental learning | **0.852** | 0.741 | **0.792** | **0.657** |

As hypothesised, the introduction of hard negative training examples substantially increased the precision of the network, from a value of 0.7327, to 0.8524. Recall dropped very slightly, likely due to proportionally less training time devoted to root locations. Despite this, Dice and UoI scores increased, with a UoI score of 0.6570 representing an overlap with the ground truth 0.2464 higher than the closest competing approach.

Figure 8 illustrates the reduction in noise attributed to hard negative training, with 3D rendered views as well as sample segmented slices. Red circles highlight areas of noise that are no longer segmented by the network following retraining.

**Figure 8:**
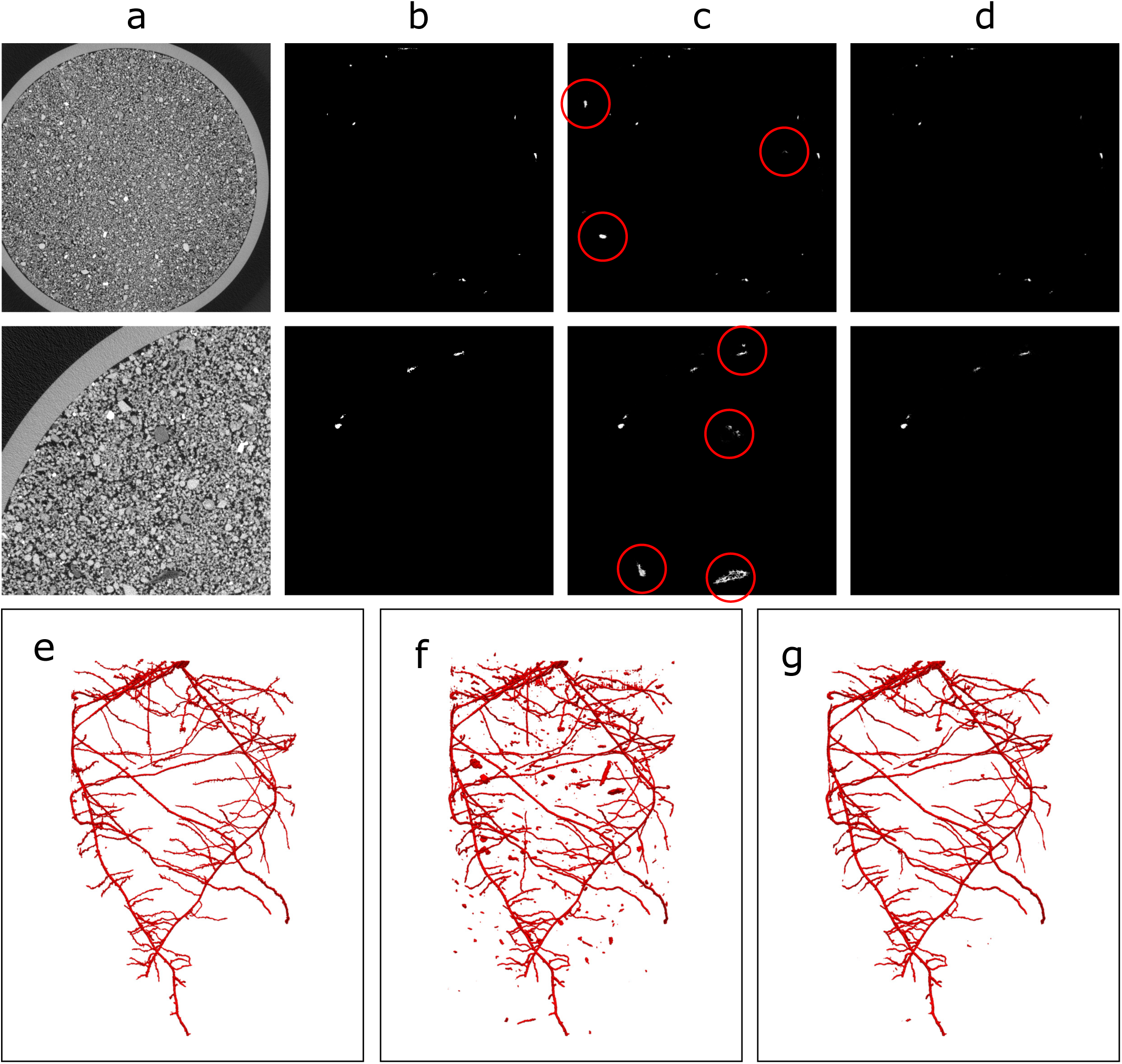
Axial view (top and middle rows) and volume rendered (bottom row) from the segmentation masks: a) original CT image, b) manual segmentation, c) MRML without incremental learning, d) MRML with incremental learning. Red circles indicate false positive segmentations. e) rendered volume of the ground truth for the plant in the top row., f) rendered volume of MRML without incremental learning, g) rendered volume of MRML with incremental learning.

## 4 DISCUSSION

The proposed MLMR network offers accuracy improvements over traditional bottom-up image analysis, as well as newer deep learning architectures. Beyond the accuracy improvements, there are a number of other key benefits in segmenting volumes using a patch-based volumetric approach. Commonly used tools such as RooTrak and Root1 make assumptions about the connectivity of root systems that patch-based deep learning does not. RooTrak uses level sets to track the movement and growth of roots moving downward through the stack. This raises complexity where roots travel sideways, or upwards, and requires the tool to spawn additional processes to handle these events. In addition, as the positions of roots in slice t are dependent on successful segmentation at slice t-1, failures cannot be recovered, while the level set’s curvature term that enforces smoothness of root edges it also disproportionately penalises fine roots with small diameter. We have also found that RooTrak often misclassifies roots travelling along the edge of the container as background, requiring manual restart. This is a failure mode that we have not observed in the MLMR architecture.

Root1 makes different assumptions about the root system, requiring additional post-processing to achieve the best results. Pixels are segmented individually based on local image information, but due to the background noise produced, a connected component algorithm is used to extract the main root system. This technique assumes that the segmentation has fully connected all roots, something that cannot be guaranteed in practice. Mathematical dilation is often used to fill small gaps, but this is a human-controlled and subjective process, and larger gaps will remain.

MLMR makes no assumptions about root system connectivity, beyond any shape constraints enforced by the learned filters. Each volumetric patch of image is considered in isolation, meaning that any failures are not propagated to nearby regions. This means that issues such as contrast changes throughout the volume are ignored, as roots are considered only against the background in the patch in which they reside. This patch-based approach has the additional advantage of making the system parallelisable, which is helpful given the computational demands of 3D imaging pipelines. Indeed, a single thread, single GPU implementation of any of the deep networks run over a large volume may take up to 10 hours to complete, a multi-threaded implementation will require a fraction of this time.

The advantage of the RooTrak and Root1 approaches is that noise — errors in which background is incorrectly segmented as root material — are explicitly removed as part of the process. RooTrak does not explore image regions far from the level set, and Root1 only preserves pixels connected to the main system. We perform no post processing, and thus do not suppress any noise should it be output in the segmentation. False positives typically occur where organic particles appear with similar size and shape to small roots. Other false positives are related to unconnected roots, perhaps remaining in the soil from another source, that are picked up using our approach. We decided to avoid any case specific post-processing in this work, and we have focused on minimising noise through careful network design and training. Nevertheless, additional techniques for identifying and removing false positives are an avenue for future work. In its current form, we found that it was possible to extract the primary root system for quantification using an existing tool [38]. This tool traverses the primary root system from detected tip locations to the seed, and fits smooth polynomial splines to major roots. The tool extracts the architecture and outputs in the widely adopted RSML format [39].

## 5 CONCLUSION

In this paper a novel multi-resolution encoder-decoder was proposed for volumetric segmentation of roots from soil in CT images. The method used volumetric images that had been previously annotated by an expert user.

The proposed network, MLMR, comprises two encoding and decoding paths, separately responsible for dealing with different image resolutions. The native path handles small patches of image at high resolution, whereas the downsampling path considers a coarse but wider field of view. Features from the wider field of view are cropped appropriately to spatially align with the native path, are concatenated, and used for segmentation by the final layers of the network. Multiple loss functions were used to train the system, ensuring that each path learned useful features that contributed to overall network performance. We also demonstrate that incremental learning through hard negative images improves the precision of the network, a process that did not add substantial time to the overall training process.

We compared our approach to an existing and popular root phenotyping tool, as well as a number of commonly used encoder-decoder networks, such as a stacked hourglass [25] and a volumetric U-Net [33]. Our results demonstrate that deep learned approaches often outperform traditional imaging techniques, but also that single resolution deep architectures often sacrifice either recall or precision, based on their configuration. Native resolution offers better segmentation of root boundaries and fine roots but increases noise. Wider fields of view reduce noise, but also under segment root material. MLMR was able to successfully segment with both high recall and high precision, resulting in overall scores on Dice and IoU above any competing technique.

All code, data and trained models will be released publicly. Future work in this area will continue to improve network architectures to provide finer segmentation detail, and to utilise transfer learning to explore the use of these networks in different imaging or scanner settings and for different species. We will also explore the use of a wider variety of soil types, with a view to establishing a richer training dataset and improving the robustness of these deep learning techniques to changes in the input. Overall, we have demonstrated that a multi-resolution encoder-decoder approach can automatically segment plant roots from CT soil images. Adoption of this tool will greatly reduce human involvement in segmentation of CT images; MLMR is therefore integral for enhancing throughput and consequently advancing knowledge of root-soil interactions and plant selection in breeding programs.

## ACKNOWLEDGMENTS

The work reported here was funded by the US Dept of Energy via the ARPA-e ROOTS project Low Cost X-Ray CT System for in-situ Imaging of Roots. C.S. and M.G. and thank the European Research Council (ERC) for FUTUREROOTS project funding. M.P. would like to thank the University of Nottingham and the Future Food Beacon of Excellence for fellowship funding that contributed towards this work.

